# Rapid and reversible optogenetic silencing of synaptic transmission by clustering of synaptic vesicles

**DOI:** 10.1101/2022.06.13.495966

**Authors:** Dennis Vettkötter, Martin Schneider, Jana Liewald, Sandra Zeiler, Julia Guldan, Shigeki Watanabe, Alexander Gottschalk

## Abstract

Silencing neurons acutely and specifically informs about their functional roles in circuits and behavior. Existing optogenetic silencers include ion pumps or channels, and tools that damage the neurotransmitter release machinery. While the former hyperpolarize the cell and can alter ionic gradients, the latter allow only slow recovery, requiring *de novo* synthesis. Thus, there is a need for other strategies combining fast activation and reversibility. Here, we use light-evoked homo-oligomerization of cryptochrome CRY2 to silence synaptic transmission, by clustering synaptic vesicles (SVs). We benchmark this tool, optoSynC, by electrophysiology and locomotion in *Caenorhabditis elegans*. optoSynC clusters SVs within 25 s, causing approximation, observable by electron microscopy. Locomotion silencing is rapid (tau_on_ ∼15 s) and recovers quickly (tau_off_ ∼10 min) after light-off. Further, optoSynC can inhibit exocytosis for several hours, at very low light intensities. optoSynC is a highly efficient, ‘non-ionic’ optogenetic silencer that may further allow to manipulate different SV pools.

## Introduction

Neurons are highly specialized cells that transmit intercellular information via electrical and chemical signals. In terminals of chemical synapses, three kinds of synaptic vesicle (SV) pools are typically distinguished: the reserve pool (RP), the readily releasable pool (RRP), and the recycling pool^1,2^. New SVs that are filled with transmitter reside in the RP. During the SV cycle, SVs can be recruited from the RP to the active zone plasma membrane. Following arrival of an action potential, Ca^2+^ enters the terminal via voltage-gated Ca^2+^ channels (VGCCs), and this triggers the fusion of SVs with the plasma membrane. SV fusion, facilitated by SNARE proteins, leads to release of neurotransmitters into the synaptic cleft^3,4^. Current models describe the processes underlying chemical synaptic transmission, at the level of the SV, by the steps of docking, priming, and fusion/exocytosis^5^. Following exocytosis, several SV-proteins have been shown to remain clustered in the plasma membrane, thus facilitating their recycling by ultrafast endocytosis. The formation of new SVs from the synaptic endosome and their refilling with neurotransmitters within the RP concludes the SV cycle^6-8^.

To study nervous system function, and the underlying molecular and cellular processes, the ability to modulate neuronal activity in a reversible way is instrumental. Ideally, methods allowing such modulation are applicable in intact animals. A major development in this context are the various techniques to influence neuronal activity with light, subsumed under the term optogenetics^9,10^. Light-responsive proteins are expressed in cells to affect their physiology in various ways. The first optogenetic protein used is channelrhodopsin-2 (ChR2), a blue light-gated cation channel^11^. When heterologously expressed, the blue light application induces depolarization of cultured mammalian neurons to trigger action potential-driven synaptic transmission. In the intact nematode *Caenorhabditis elegans* (*C. elegans*), even certain behaviors are induced^12,13^. Since then, many labs have adapted the tool to interrogate the circuit-behavior relationships in several model organisms. Importantly, by turning off the illumination, the neuronal excitation is rapidly terminated (typically within 20-100 ms), thus ChR2-based optogenetics is generally highly reversible.

Besides neuronal excitation, inhibition of neural activity is highly informative about the function of neurons within circuits. Several silencing strategies using genetically encoded tools with diverse biophysical properties were developed over the past ca. 15 years^14^. Light-activated ion pumps and channels enable rapid and reversible hyperpolarization of neurons, thus resulting in suppression of neuronal activity with high spatial and temporal resolution^15-18^. However, these tools still exhibit a decline in efficacy when stimulated over a prolonged period^19^, for two reasons: First, while being highly efficient tools for silencing, the inhibitory action of anion channelrhodopsins (ACRs) depends on membrane potential and the chloride gradient. Also, they can affect intracellular chloride concentration and alter cellular physiology, and in synaptic terminals, they may even cause depolarization instead of hyperpolarization^20,21^. Other silencing techniques, such as G-protein-coupled receptors (GPCRs), are limited in the often ill-defined coupling specificity to heterotrimeric G proteins. GPCR signaling is of slow onset kinetics (dozens of ms to sec), and thus, GPCRs are mostly used in applications that do not require fast activation^14^. Silencing by excitation, an approach utilizing the expression of ChR2 in inhibitory interneurons has the limitation that specific subtypes of neurons cannot be targeted precisely^14,22^.

Many of these inhibitory tools are ill-suited for long-term silencing of neuronal activity. Therefore, other tools have been developed to damage or degrade proteins essential for synaptic release, like miniSOG (miniature singlet oxygen generator), which upon blue-light stimulation generates reactive oxygen species^23-26^. These radicals oxidize susceptible residues on nearby proteins, such as cysteine, histidine, methionine, tryptophan, and tyrosine. When attached to SV proteins like the SNARE synaptobrevin (VAMP2) or synaptophysin (SYP1), application of blue light for short periods (seconds to minutes) leads to the inactivation of the SNARE complex^25^. This approach is reversible, however, only at a slow time scale, as it requires the *de novo* synthesis of the affected proteins. Also, the generation of damaging radicals is likely to have off-target effects, on other synaptic proteins in the proximity of the target proteins like SNAREs, and also proteins in the secretory pathway. Therefore, longer-lasting effects on synaptic strength and neuronal cell biology are conceivable but not well understood. Along the same lines, yet aiming to avoid off-target effects, a photosensitive degron (PSD) has been adapted for applications in the nervous system. The PSD enables the degradation of specific proteins, triggered by light^27^. Fused to synaptic proteins, it enables higher precision for targeting of the damaging effects^26^. Light-induced degradation of synaptotagmin (SNT-1), required for synchronous neurotransmitter release, resulted in inhibition of neurotransmission to an extent comparable to miniSOG. However, this approach works only in the absence of endogenous SNT-1 (i.e. null mutants), or the PSD must be inserted in the genomic locus. Another approach targeting the SNARE complex is photoactivatable botulinum neurotoxin (PA-BoNT)^28^, which cleaves VAMP2 in a light-dependent manner. PA-BoNT does not require constant illumination for long-lasting effects. Yet, as for miniSOG or PSD, reversibility takes up to 24 h, as damaged proteins must be synthesized *de novo*. While miniSOG takes some minutes for full effect, PSD and PA-BoNT are effective after 30-60 min stimulation, and thus, the onset is rather slow.

Therefore, there is still a high demand for silencing tools that work with high spatial and temporal precision and have fast onset and recovery kinetics with sustained silencing qualities. Since the SV cycle and chemical synaptic transmission require the mobilization and approaching of SVs to the active zone membrane, sequestering of SVs may inhibit synaptic transmission. To achieve this, one may use cryptochrome-2 (CRY2), which can undergo light-dependent di- or oligomerization^29^. CRY2, derived from the plant *Arabidopsis thaliana*, is the most widely used cryptochrome in optobiological studies. Stimulation of CRY2 with blue light (∼450 nm) induces protein-protein interactions like homo-oligomerization^29,30^ and heterodimerization with the cryptochrome-interacting basic helix-loop-helix protein 1 (CIB1)^31,32^. These interactions were mapped to the photolyase homology region (PHR), containing the chromophore FAD. Compared to other tools, such as iLID^33^ or Magnets^34^, also CRY2 can act as a light-inducible dimer, however, it can also act as a single-component system. Therefore, only one protein has to be introduced into the host organism. Previous studies utilized CRY2 heteromerization to trap target proteins into complexes using light^35^. This ‘light-activated reversible inhibition by assembled trap’ (LARIAT) was recently used to interfere with synaptic transmission by targeting VAMP2^36^. These authors suggested that their approach inhibits synaptic transmission by blocking the SV release machinery, i.e. somewhat similar to the PSD and miniSOG approaches. However, using the homo-oligomerizing properties of CRY2 may enable cross-linking of SVs already before they reach the plasma membrane, thus inhibiting synaptic transmission. Furthermore, such an approach may allow investigation of various aspects of the SV cycle, if SV mobilization from the RP or their transport to the active zone is inhibited.

Here, we used a variant of CRY2, CRY2olig(535), that combines the homo-oligomerization inducing mutation E490G, and a truncation that reduces dark activity, and fused it to the SV protein synaptogyrin (SNG-1), a tetraspan vesicle membrane protein^37^. This yielded optoSynC, an <u>opto</u>genetic tool for <u>syn</u>aptic vesicle <u>c</u>lustering. By behavioral and electrophysiological assays, we show that optoSynC can efficiently inhibit synaptic transmission in different subtypes of neurons within seconds, allows for long-term silencing for several hours, and can recover in the absence of light within minutes. By electron microscopy, we demonstrate that SVs show a marked clustering, in response to light activation, thereby indicating that synaptic transmission is blocked, at least in part, by affecting SV mobility.

## Results

### Development of optoSynC, an optogenetic tool for clustering of synaptic vesicles

Synaptogyrins are highly abundant SV proteins^38^. However, loss-of-function mutations of *C. elegans sng-1* neither result in major defects in synaptogenesis nor neuronal activity^37^. Possible functions in clathrin-independent endocytosis were observed only along with other mutations^39^. Therefore, SNG-1 can serve as an ‘inert’ anchor to attach proteins to the SV, as we did previously for PA-BoNT^28^.

To interfere with synaptic transmission by SV clustering, we first tried a LARIAT-based approach, utilizing the heterodimerization of CRY2 with its interaction partner CIB1. Enhanced CRY2 variants with positive charges at the C-terminal residues, i.e. E490G, promote light-induced homo-oligomerization; this tool was termed CRY2olig^40^. To improve expression and to reduce self-association in the dark, a truncated CRY2 module, CRY2(535), was developed^41^. We fused CIB1, tagged with GFP, to SNG-1 and expressed an mOrange2-tagged CRY2olig(535) in the cytosol of neurons. Our design resembled the recently reported opto-vTrap^36^. We hypothesized that blue light illumination (470 nm) should activate the PHR domain of CRY2 (Ref. ^42-44^), inducing dimerization, and may thus trap SVs in clusters and inhibit neurotransmission (**Supplementary Fig. 1a**). To explore this, we assayed locomotion behavior (swimming), as this is very sensitive to malfunction of the motor neurons. However, illumination did not alter swimming behavior (**Supplementary Fig. 1b, c**). Thus, the LARIAT approach appears not to work in *C. elegans*.

We therefore turned to the homo-oligomerization properties of CRY2^45^. Though this mechanism is complex and only partially characterized^46^, many CRY2 optogenetic tools rely on it^30,40,47^. We fused CRY2olig(535) to the C-terminus of SNG-1 and introduced the construct, hereafter termed ‘optoSynC’ (<u>opto</u>genetic <u>syn</u>aptic vesicle <u>c</u>lustering), in *sng-1(ok234)* null mutants, expressing it from a pan-neuronal promoter. We examined if light application has any effects, e.g. induce oligomerization, thus trapping SVs in clusters (**Fig.1a**), or blocking release sites in the plasma membrane (not shown) and perturbing neuronal activity. To this end, we recorded swimming cycles under dark and light conditions (**Fig. 1b, c**). Activation of optoSynC (470 nm, 0.1 mW/mm^2^, 5 s) led to a drastic slowing of swimming behavior (Supplementary Video 1). Swimming cycles decreased by 80% within the first 25 s after one light pulse (τ = 14.18 s; **Fig. 1b**). Application of further light pulses in a 5 s / 25 s inter-stimulus interval maintained the inhibition of swimming cycles for minutes. After the end of the illumination, animals recovered normal swimming behavior within ca. 20 min (**Fig. 1c**, plateau following one-phase association, τ = 9.79 min). This indicates that the optoSynC construct returned to the dark state, possibly releasing SVs from the putative clusters.

**Figure 1.**
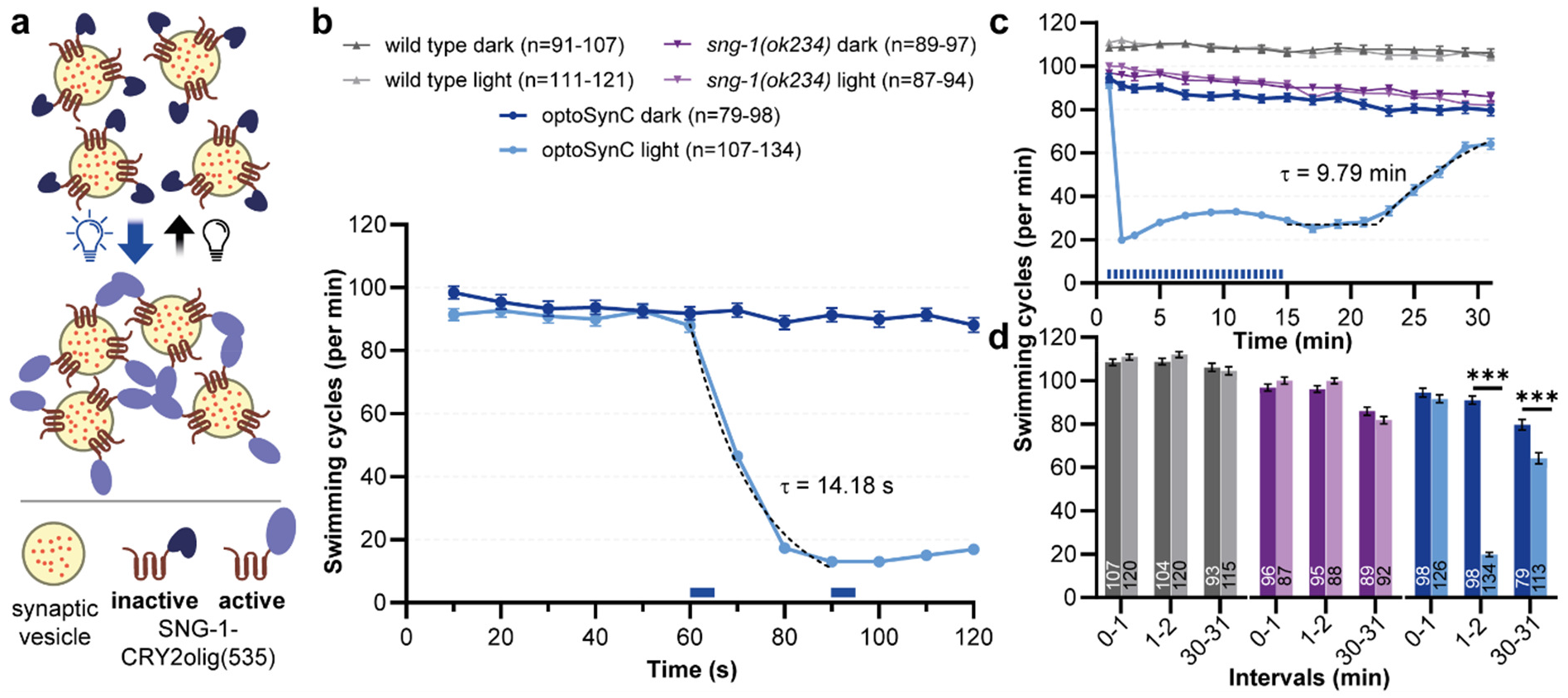
OptoSynC inhibits behavior within seconds and recovers within minutes in the dark: **a** Schematic illustrating SV clustering through homo-oligomerization of CRY2olig(535) upon blue light illumination. **b** Mean (± s.e.m.) swimming cycles of worms expressing optoSynC pan-neuronally. Blue light (470 nm, 0.1 mW/mm^2^, 5 s/ 25 s ISI) is indicated by blue rectangles; dotted line: one-phase decay fit (60 s - 90 s). **c** As in (b), extended time course. Sustained inhibition of swimming by ongoing light pulses, and recovery in the dark. Dotted line: ‘plateau followed by one phase association’-fit. **d** Group data before (0-1 min), during (1-2 min) and after (30-31 min) of blue light illumination. Two-way ANOVA with Bonferroni correction between light and dark measurements of wild type, *sng-1(ok234)* and optoSynC expressing animals (*sng-1* background); *p<0.05, **p<0.01, ***p<0.001. Number of animals is indicated in each bar.

To facilitate visualization of the SV cloud via optoSynC, we inserted a fluorescent protein between SNG-1 and CRY2olig(535). However, this reduced the functionality of optoSynC by 50% (**Supplementary Fig. 2a-c**). Further, we characterized optoSynC efficacy under different illumination protocols and light intensities. Using continuous vs. pulsed blue light did not increase the effect on swimming behavior (**Supplementary Fig. 2d**). When we reduced the light intensity from 0.1 mW/mm^2^ to 1.4 µW/mm^2^, more light pulses were necessary to achieve a strong effect. However, swimming could still be reduced by approximately 65% when 4 pulses (two per minute) where applied (**Supplementary Fig. 2e**). In sum, optoSynC is a highly sensitive tool that inhibits locomotion within 25 s after light-stimulation, likely by affecting synaptic transmission, and recovers within 20 min in the dark.

### OptoSynC can alter behavior instantaneously

OptoSynC activation strongly inhibits locomotion, likely by blocking SV mobility and/or fusion. Since analysis of swimming behavior requires at least 10-20 seconds of data, it is not suited to precisely determine how fast optoSynC may act. We thus also analyzed crawling locomotion speed. The short-wavelength light receptor LITE-1 mediates an escape response^48^, which results in an increased crawling speed upon illumination. We analyzed the crawling speed of animals expressing optoSynC; wt and *sng-1(ok234)* were used as controls. Transgenic animals are in *sng-1(ok234)* background, unless otherwise stated. To evoke the escape response, we illuminated animals with blue light pulses (5 s/ 25 s ISI, 1 mW/mm^2^; this light intensity is more than required to activate optoSynC; however, it is used to evoke robust escape behavior). Upon illumination, crawling speed of wt and *sng-1(ok234)* animals rapidly increased by approximately 100% within 5 s, and decayed again after the light was turned off. However, transgenic animals, as blue light simultaneously activates optoSynC, were unable to accelerate as much as the controls (**Fig. 2a-c**). Already the behavioral response to the first light pulse was largely attenuated, and speed increased by only ca. 20%. For subsequent stimuli (5 s / 25 s ISI), almost no increase was observable, while non-transgenic animals always markedly increased their speed. Thus, optoSynC inhibits the escape response instantly, and the time course of 25 s observed during swimming (**Fig. 1b**) is rather due to the analysis method, and neuronal silencing during swimming is likely similarly fast. To control for the effect of blue light, we used *lite-1(ce314)* animals. These animals are unable to detect blue light, and thus do not show escape behavior. optoSynC activation did not significantly decrease the basal crawling speed (**Fig. 2c**). As an additional control, we mutated the FAD-binding pocket of CRY2 at position D387A, which renders CRY2 photoinactive^32^. Compared to *sng-1(ok234)* animals, animals expressing the resulting optoSynC-DA responded similarly to the blue light pulses (**Fig. 2a-c**). Thus, expression of optoSynC does not *per se* affect synaptic transmission, unless it is photo-activated.

**Figure 2.**
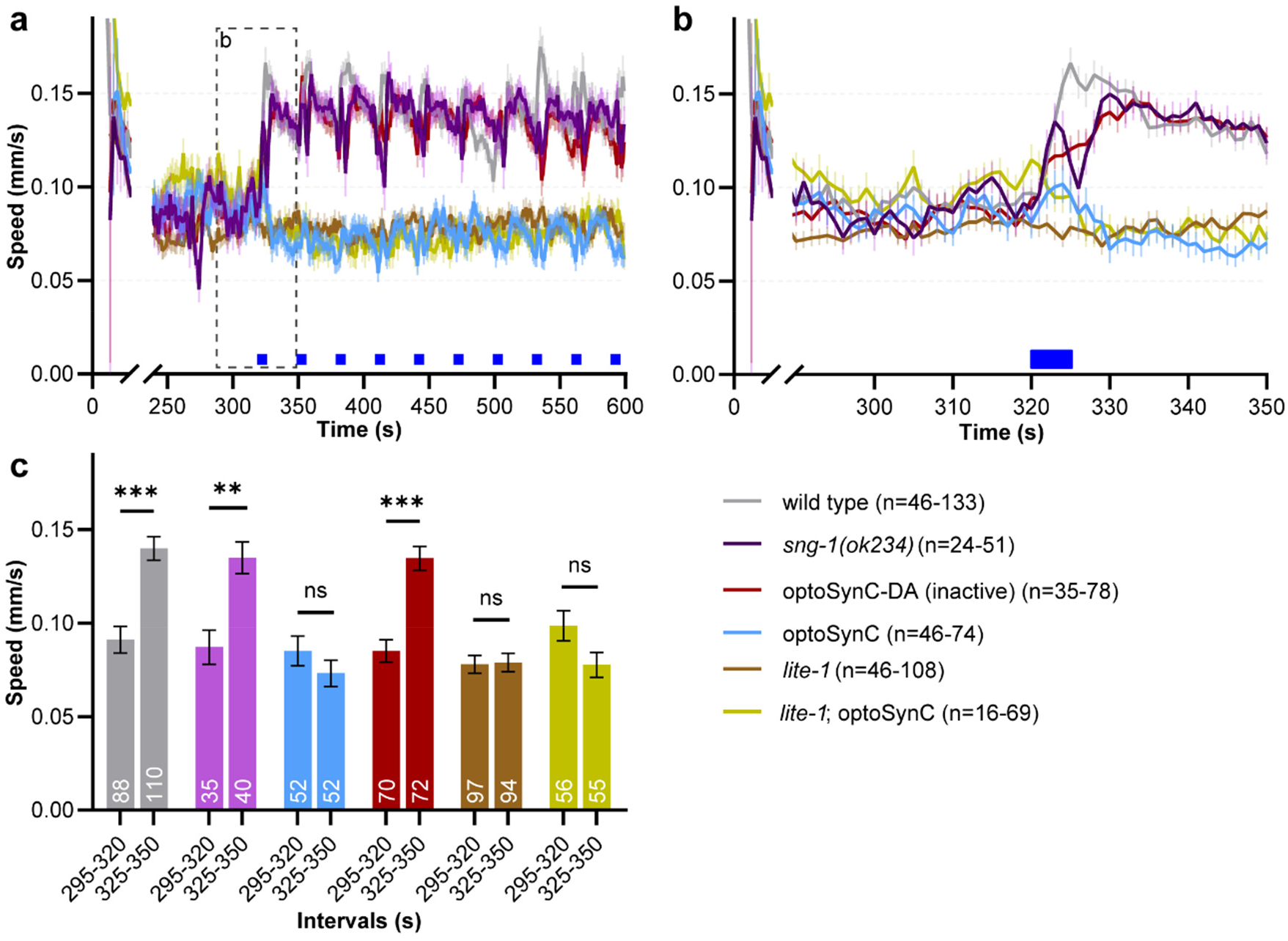
Immediate inhibition of the LITE-1-dependent escape response by optoSynC: **a** Crawling speed analysis, blue light application (470 nm, 1 mW/mm^2^, 5 s / 25 s ISI, indicated by blue rectangles), genotypes as indicated, all animals in *sng-1(ok234)*, unless stated. **b** Close-up of box indicated in a; mean ± s.e.m. **c** Group data, mean (± s.e.m) crawling speed analyzed in time intervals before (295-320 s), and after (325-350 s) first light pulse. Two-way ANOVA with Bonferroni correction: *p<0.05, **p<0.01, ***p<0.001, ns not significant. Number of animals shown in the bars.

### optoSynC activation strongly decreases mPSC frequency and can reduce cholinergic transmission for hours

To more directly examine the effect of activated optoSynC on SV release, we recorded miniature post-synaptic currents (mPSCs) from muscle cells, which are innervated by motor neurons expressing optoSynC. mPSC frequency was significantly reduced in response to optoSynC activation (0-30 s: 33.29 ± 4.74 s^-1^; 95-105 s: 13.35 ± 2.27 s^-1^), as compared to wt (0-30 s: 33.05 ± 6.09 s^-1^; 95-105 s: 34.84 ± 6.77 s^-1^; **Fig. 3a-d**). This indicates a rapid and robust decrease of SV fusion events upon activation of optoSynC, possibly by trapping of SVs in vesicle clusters. mPSC amplitude remained unchanged upon illumination (**Fig. 3e, f**). As the mPSC amplitude is determined by loading of SVs with neurotransmitters and/or SV size^49^, these results suggest that optoSynC has no effect on SV properties.

**Figure 3.**
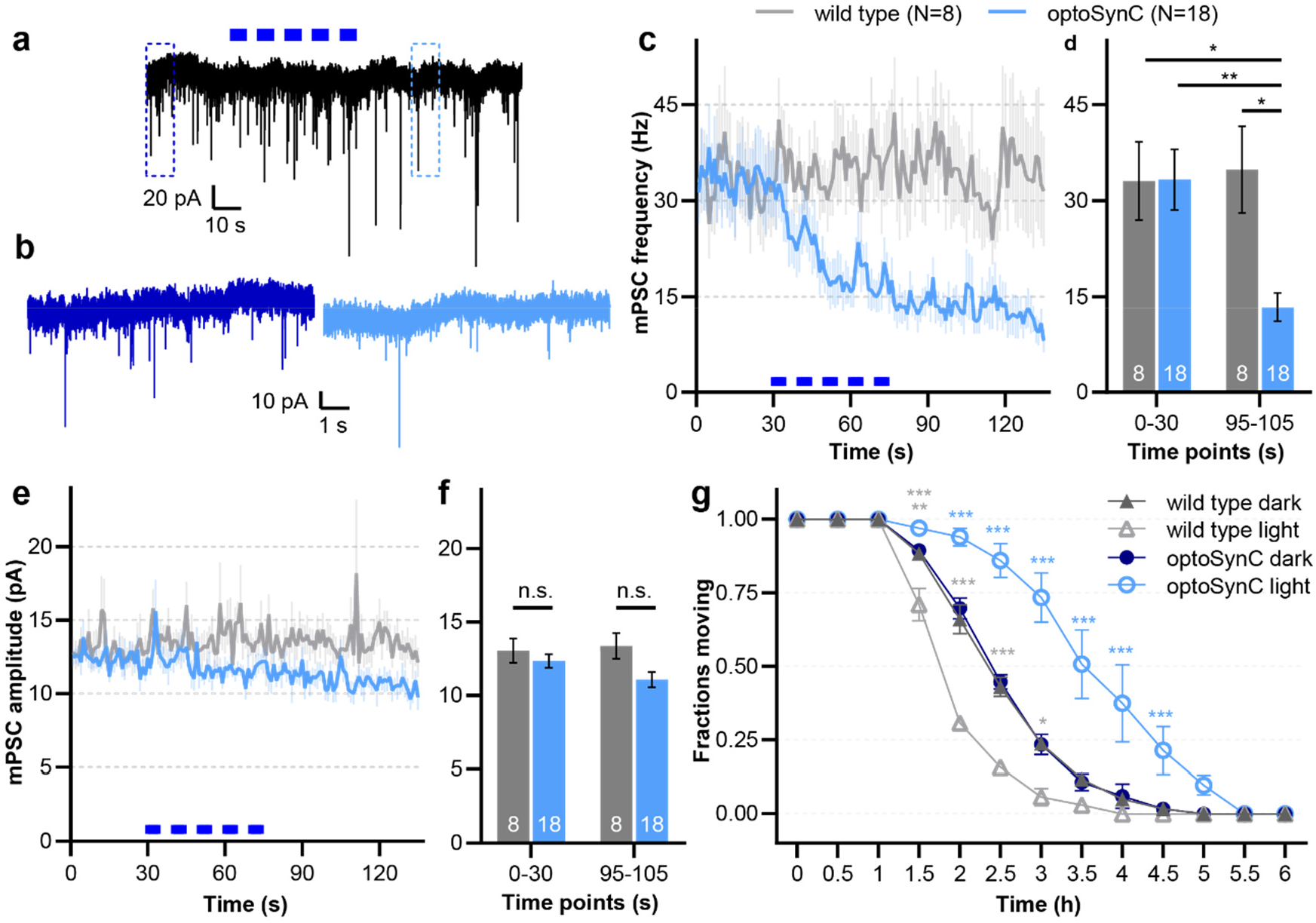
optoSynC stimulation reduces miniature post-synaptic current (mPSC) rate at the neuromuscular junction (NMJ) and can block cholinergic transmission for hours: **a** Representative postsynaptic current traces recorded in body wall muscle cells of optoSynC-expressing animals; blue light pulses (470 nm, 8 mW/mm^2^, 5 s / 5 s ISI) indicated by blue rectangles. **b** Close-up of regions indicated in (a), before (0-10 s, dark blue) and after (95-105 s, light blue) blue light illumination. **c** Mean ± s.e.m. mPSC frequency with blue light illumination of wild type and optoSynC expressing animals. **d** Group analysis of data in (c), intervals before (0-30 s) and after (95-105 s) illumination. Two-way ANOVA with Bonferroni correction; *p<0.05, **p<0.01, ***p<0.001). Number of animals is indicated. **e, f** Analysis of mPSC amplitude, as in (d, e). **g** Paralysis of animals in response to 2 mM aldicarb under continuous blue light illumination (470 nm, 0.05 mW/mm^2^). Two-way ANOVA with Bonferroni correction of N=3 experiments averaged at the indicated time points with n=15-27 animals each, *p<0.05, **p<0.01, ***p<0.001; error bars are s.e.m.

Patch-clamp recording of *C. elegans* muscles requires dissection of the animal, and the preparation can be recorded reliably only for a few minutes^50^. Therefore the approach is not suited for analysis of long-term effects of optoSynC on cholinergic transmission and its potential for long-term inhibition. To this end, we used a pharmacological assay in intact animals. Incubating animals in the acetylcholinesterase inhibitor aldicarb causes accumulation of acetylcholine (ACh) in the synaptic cleft, thus leading to progressive paralysis^51^. We compared animals with and without the pan-neuronal optoSynC, permanently stimulated with low-intensity light (470 nm, 0.05 mW/mm^2^) or kept in the dark (**Fig. 3g**). Animals kept in darkness throughout the experiment were paralyzed at the same rate as non-transgenic controls. Blue light stimulated animals expressing optoSynC paralyzed significantly more slowly than controls, most likely due to inhibition of ACh release. In contrast to optoSynC expressing animals, wt animals stimulated with blue light paralyzed significantly faster than dark controls. Conceivably, this is due to the evoked escape response mediated by activation of LITE-1 by blue light, which causes increased cholinergic transmission (**Fig. 2**).

### Ultrastructural analysis in optoSynC synapses unravels light-induced SV clustering

The behavioral and physiological effects of optoSynC activation could result from clustering of SVs, which may thus not be mobilized from the RP. Alternatively, proteins of the SV, particularly oligomerized SNG-1::CRY2olig(535), may remain in the active zone membrane, instead of being recycled by ultrafast endocytosis^52-56^. This may prevent SV recycling or docking and priming of further SVs during ongoing stimulation. Thus, to examine the mode of action of optoSynC in detail, we analyzed the ultrastructure of cholinergic synapses expressing optoSynC by serial section transmission electron microscopy (TEM)^53^. Animals were illuminated for 5 s (+light; 470 nm, 0.1 mW/mm^2^) and high-pressure frozen (HPF), 25 s following the pulse, to ensure maximal inhibition of synaptic transmission (**Fig. 1b, Supplementary Fig. 2d**). Control animals (-light) were kept in darkness during the HPF process. HPF samples were freeze-substituted, stained, and 40 nm thin sections were analyzed. Plasma membrane (PM), dense projection (DP), cytosolic SVs, docked / tethered SVs, dense core vesicles (DCVs), and large vesicles (LVs) were annotated as described in Ref. 57 (**Fig. 4a, b, Supplementary Fig. 3a, b**). To explore whether SVs became clustered, we quantified the distance of each SV to its nearest neighboring SV in the same micrograph (**Fig. 4c, Supplementary Fig. 3c, d**). Photostimulated synapses showed significantly smaller nearest distances (42.30-20.28 nm, 26.55 nm median) when compared to unstimulated controls (50.82-23.35 nm, 31.01 nm median); **Fig. 4c, d**). To exclude that these effects were overestimated by analyzing individual SVs, we also calculated the mean of the nearest distances per micrograph (+light: 45.6-31.45 nm, 37.20 nm median; -light: 54.94-40.32 nm, 47.10 nm median; **Fig. 4e**), and per synapse (i.e. in consecutive sections; +light: 40.65-33.98 nm, 39.24 nm median; -light: 51.85-45.42 nm, 47.81 nm median; **Fig. 4f**). Both analyses confirmed that there is a significant decrease of distance between SVs in the photostimulated samples, arguing that SVs became markedly more clustered by optoSynC. This must occur in addition to SV clustering in the RP, which is mediated by synapsin and the actin cytoskeleton^58^. So far, we pooled the data for all SVs, i.e. those in the RP (cytosolic) and those SVs tethered and docked to the PM. When we restricted our analysis to the RP, we observed the same significant decrease of nearest SV distances following optoSynC activation (**Supplementary Fig. 3c-f**), while this did not affect the distances of docked SVs (**Supplementary Fig. 3g-j**). We conclude that optoSynC affects clustering of SVs.

**Figure 4.**
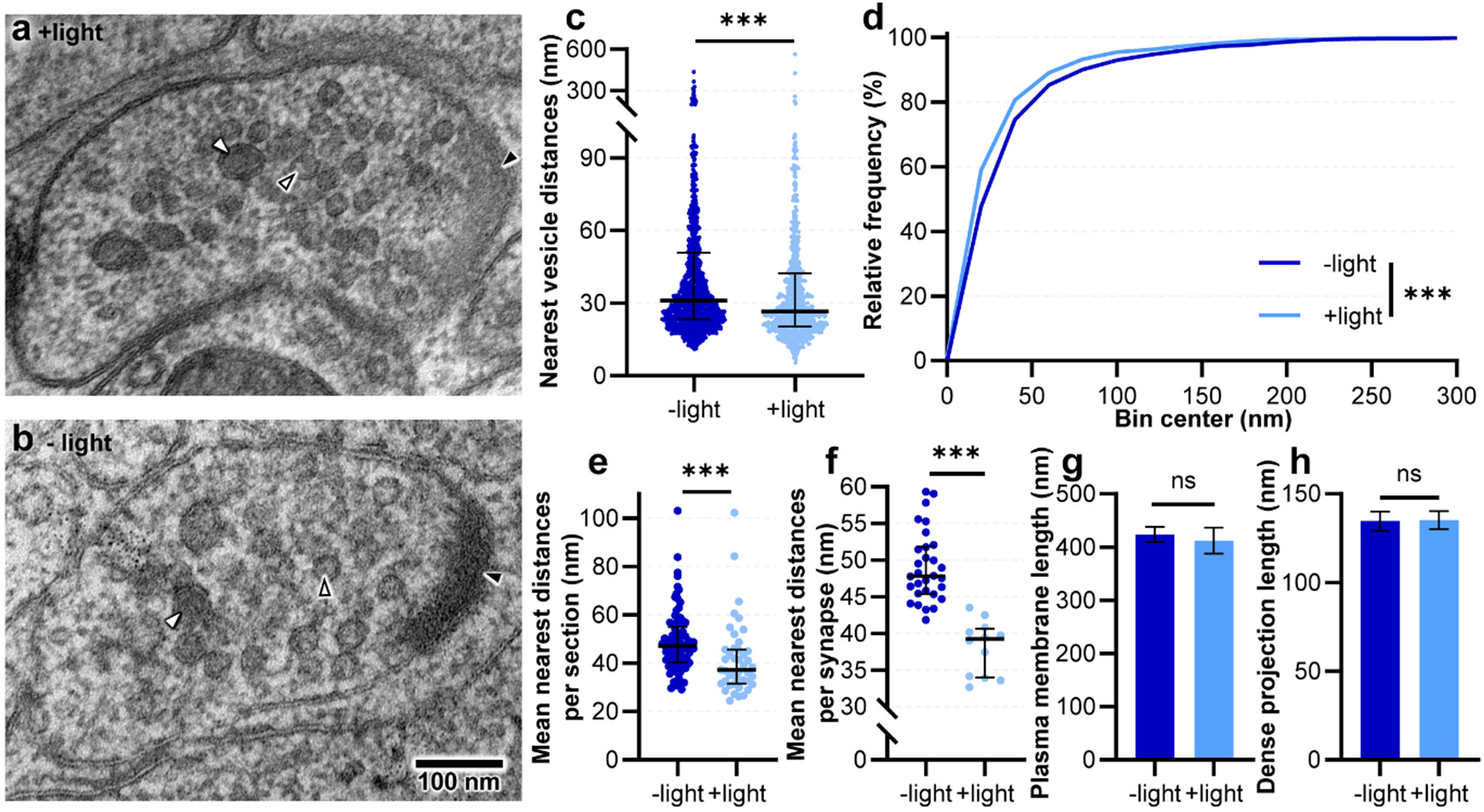
Comparing the ultrastructure of stimulated and unstimulated cholinergic synapses expressing optoSynC: **a, b** Transmission electron micrographs of representative cholinergic synapses from animals illuminated for 5 s (a, +light) with blue light (470 nm, 0.1 mW/mm^2^), or kept in darkness before (b, -light) high-pressure freezing. SVs (open black arrowheads), dense core vesicles (DCVs, white closed arrowheads) and dense projection (DP, closed black arrowheads) are indicated. **c** Distance analysis of nearest vesicles for each analyzed cholinergic micrograph shown as median with interquartile range (Mann-Whitney test between -light (n=1473) and+light (n=819), *p<0.05, **p<0.01, ***p<0.001). **d** Relative frequency distribution of nearest vesicle distances shown in c (Kolmogorov-Smirnov test, *p<0.05, **p<0.01, ***p<0.001). **e** Mean nearest distances per section shown as median with interquartile range (Mann-Whitney test between -light (n=88) and +light (n=43), *p<0.05, **p<0.01, ***p<0.001). **f** Mean nearest distances per synapse shown as median with interquartile range (Mann-Whitney test between -light (n=30) and +light (n=12), *p<0.05, **p<0.01, ***p<0.001). **g, h** Lengths of the plasma membrane (g) and dense projection (h). Mann-Whitney test between -light (n=88) and +light (n=43), *p<0.05, **p<0.01, ***p<0.001, ns not significant; Error bars are s.e.m.

Might stimulation of optoSynC also have effects on synaptic properties due to protein aggregation in the SV or plasma membrane? To this end we assessed features such as circumference of the PM (as a proxy for synapse size), length of the DP, SV diameter, or total SV number per section. No differences of PM perimeter or DP length were observed in photo-stimulated synapses (**Fig. 4g, h**). However, the diameters of SVs (+light: 36.46-20.61 nm, 38.25 nm median; -light: 33.40-29.25 nm, 31.18 nm median) and DCVs (+light: 52.37-44.45 nm, 49.43 nm median; -light: 49.62-42.67 nm, 46.37 nm median) were significantly larger after stimulation (**Supplementary Fig. 4a, b**). Our electrophysiology data did not indicate larger SV content, though (**Fig. 3e, f**); possibly, optoSynC clustering can also occur within single SVs and alter the TEM appearance of the SV membrane. Whether SNG-1 is part of DCVs, is not known. The size of LVs remained unaffected by photo-stimulation (**Supplementary Fig. 4c**), and the overall number of SVs (+light: 23.89 ± 2.83 nm; -light: 20.53 ± 1.41 nm), as well as the total number of docked SVs (+light: 1.11 ± 0.16 nm; -light: 0.93 ± 0.11 nm) was not significantly increased (**Supplementary Fig. 4d, e**). Yet, when we analyzed the distribution of docked SVs relative to the DP (**Supplementary Fig. 4f-h**), they showed a tendency of being depleted after opto-SynC stimulation. However, this excluded those SVs right next to the DP, which were significantly increased after photostimulation (**Supplementary Fig. 4f, g**). Possibly, during the time after illumination and before freezing the samples, SVs that were already docked could fuse, and as SV replenishment from the RP stopped during this time, this caused the depletion. At the DP some ‘leftover’, un-clustered SVs still docked, but then became trapped, instead of diffusing laterally into the active zone membrane^53^. Our results suggest that optoSynC blocks transmission by impeding SV replenishment from the RP, and that no major structural abnormalities are induced at the PM. However, we cannot exclude that mobility of SVs along the membrane could become restricted by optoSynC activation.

### Cell-specific expression of optoSynC allows inhibiting transmission in specific neurons

Thus far, we used pan-neuronal expression of optoSynC. Next, we asked if optoSynC also allows for inhibition of synaptic transmission in distinct neuron classes, or even individual cells. Therefore, we expressed optoSynC specifically in subsets of motor neurons, using promoters for cholinergic (p*unc-17*, encoding the vesicular acetylcholine transporter; **Fig. 5a, b**) and GABAergic neurons (p*unc-47*, vesicular GABA transporter; **Fig. 5c, d**)^59,60^. Expression and activation of optoSynC in cholinergic neurons significantly reduced swimming cycles during light-stimulation (470 nm, 0.1 mW/mm^2^, 5 s, 25 s ISI) which recovered within 20 min after switching off the stimulation (**Fig. 5a, b**). These effects were similar as for pan-neuronal expression; however, the animals did not slow down as much (reduction by ca. 55%, compared to ca. 80% for pan-neuronal optoSynC). Inhibition of GABAergic neurons by optoSynC was not as severe as for cholinergic neurons, though the reduction of swimming cycles was still significant (**Fig. 5c, d**). This is not unexpected, as GABAergic transmission is not essential for forward locomotion. We wondered if simultaneous expression of optoSynC in, and inhibition of, cholinergic and GABAergic neurons would show additive effects. However, swimming cycles were not any further reduced (**Supplementary Fig. 5e, f**). Thus, it is likely that the maximal effects we observed by pan-neuronal expression correspond to additive effects of motor neurons and upstream premotor interneurons. For highest efficiency, optoSynC should be expressed in *sng-1(ok234)* background, such that the proteins do not compete with endogenous SNG-1 for incorporation into SV membranes (**Supplementary Fig. 5a, d**).

**Figure 5.**
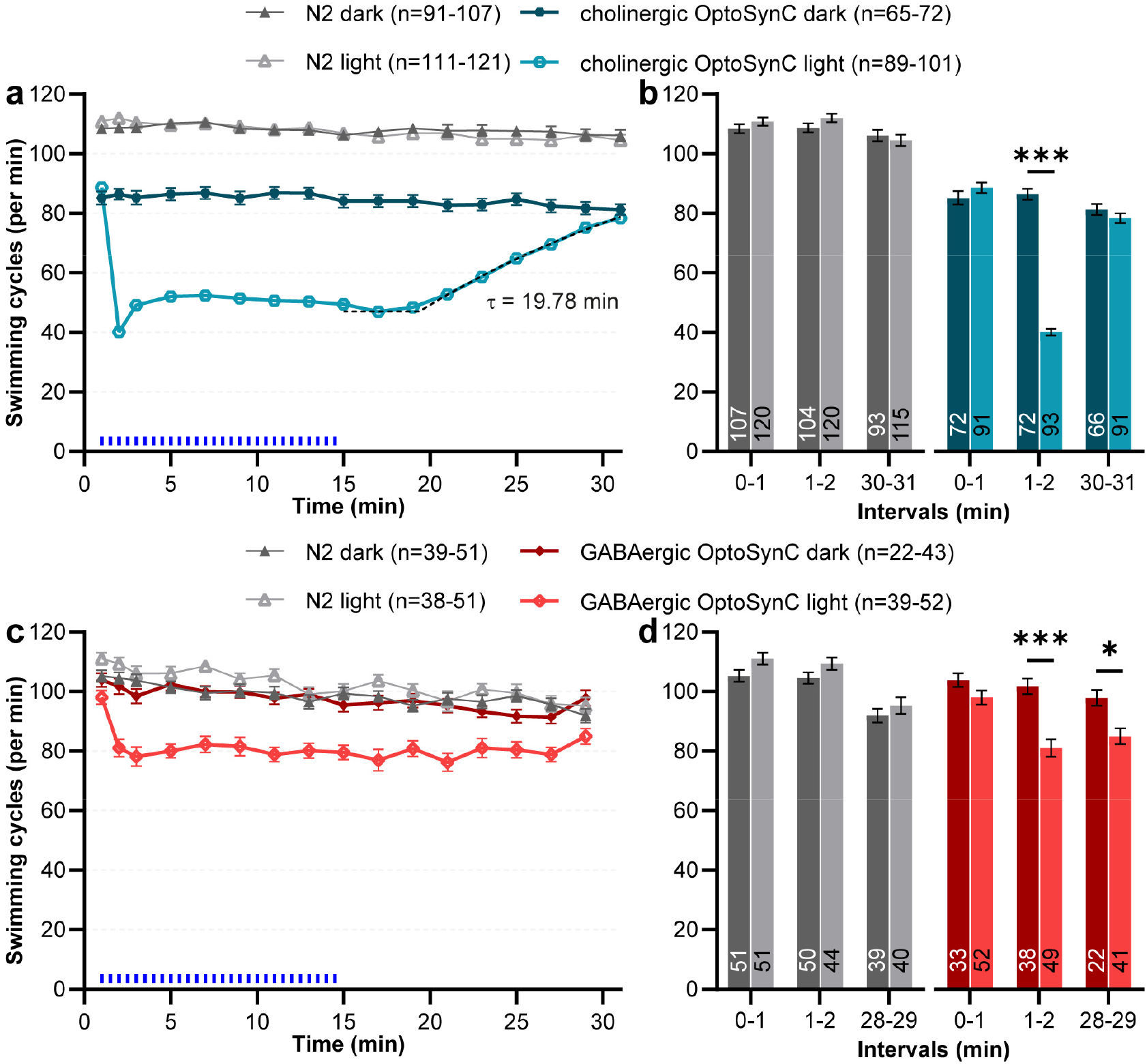
Cell-specific optoSynC inhibition of cholinergic and GABAergic neurons. **a** Swimming behavior in animals expressing optoSynC in cholinergic neurons. Blue light activation (470 nm, 0.1 mW/mm^2^, 5 s / 25 s ISI) indicated by blue rectangles. Dotted line represents ‘plateau followed by one phase association’-fit. Error bars are s.e.m. **b** Mean ± s.e.m. swimming cycles during intervals before (0-1 min), during (1-2 min) and after (30-31 min) blue light illumination. Two-way ANOVA with Bonferroni correction between light and dark measurements of wild type and cholinergic optoSynC expressing strains; *p<0.05, **p<0.01, ***p<0.001. **c, d** as for (a, b), but optoSynC was expressed in GABAergic neurons.

### Activated optoSynC blocks transmission from the single nociceptor neuron PVD

Finally, we explored whether effects of optoSynC can be observed in a single neuron and interfere with its synaptic transmission. To this end, we expressed optoSynC along with Chrimson^61^, a red-light activated channelrhodopsin, in the left/right pair of nociceptive PVD neurons^62^. Activation of Chrimson induces a rapid forward escape behavior^10^ and thus a strong increase in crawling speed (**Fig. 6a**). We thus wondered whether concomitant activation of optoSynC would cause an attenuation of this escape behavior. We compared animals expressing Chrimson and optoSynC in PVD to animals expressing only Chrimson, before and after applying a blue light pulse (5 s, 0.1 mW/mm^2^), and prior to red light stimulation (680 nm, 0.1 mW/mm^2^, 1 s / 5 s ISI). Though Chrimson is primarily activated by red light, it also shows some response to blue light^61^. Therefore, some pre-activation of Chrimson occurred with the blue light pulse used to trigger optoSynC (**Fig. 6a, b**). However, this pre-activation did not affect the consecutive activation by red light (**Fig. 6c**), and thus the velocity increase of animals expressing optoSynC and Chrimson was significantly inhibited when compared to the controls without optoSynC, or without blue light (**Fig. 6c**). In sum, optoSynC is a highly sensitive optogenetic tool for spatial, temporal, and cell-type-specific inhibition of synaptic transmission with fast onset and recovery times. It may further enable studies of subcellular SV distribution.

**Figure 6.**
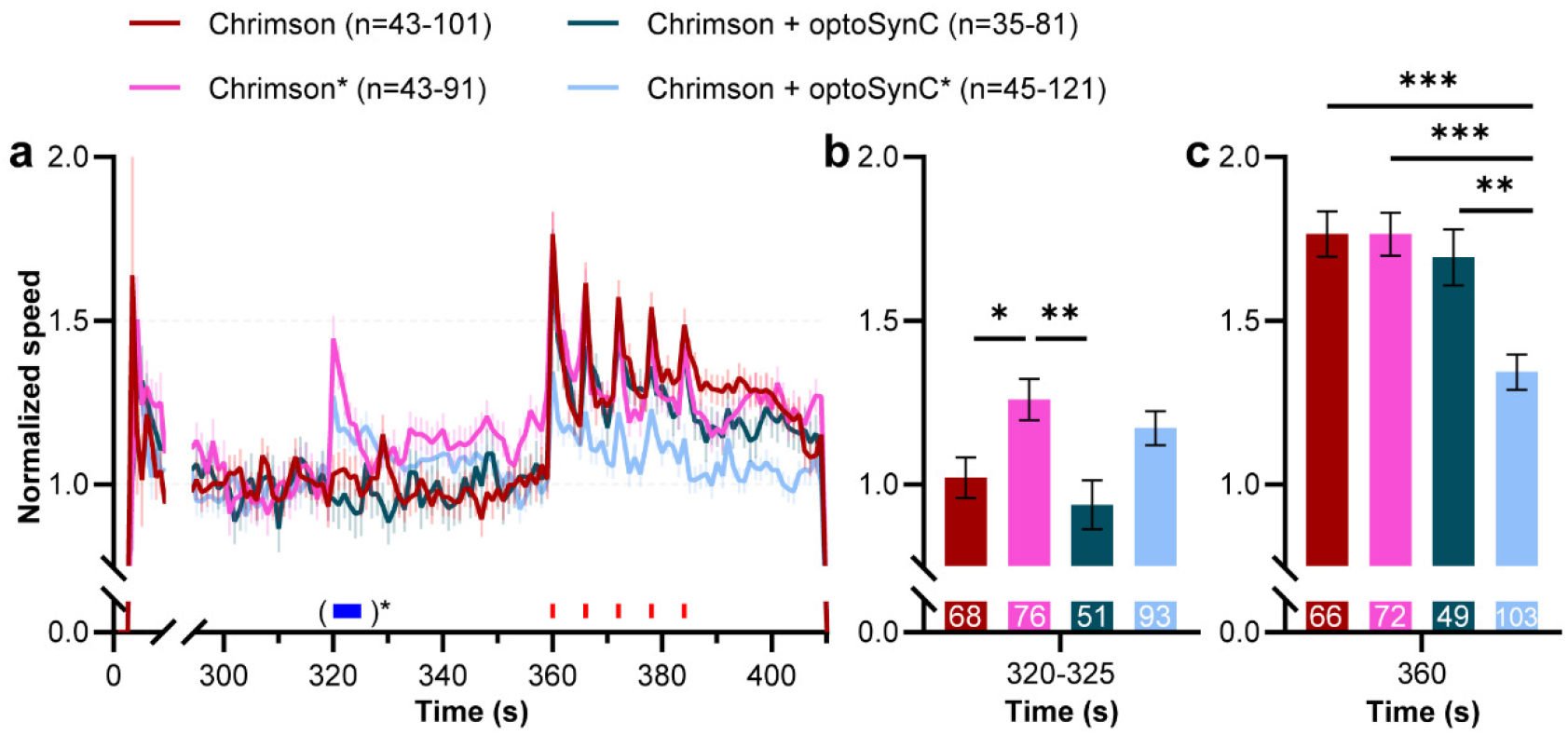
Expression of optoSynC in the single nociceptive neuron PVD attenuates PVD::Chrimson-evoked velocity increase. **a** Mean ± s.e.m. normalized crawling speed of animals expressing Chrimson, or Chrimson and optoSynC in the PVD neuron. Red light stimulation (680 nm, 0.1 mW/mm^2^, 1 s / 5 s ISI) is indicated by red rectangles, activation of optoSynC with blue light (470 nm, 0.1 mW/mm^2^, 5 s) by a blue rectangle and asterisk. **b** Mean ± s.e.m. crawling speed during time with (or without) blue pulse (320-325 s). One-way ANOVA with Bonferroni correction; *p<0.05, **p<0.01, ***p<0.001. Number of animals is indicated. **c** As in (b), but during first red light pulse (360 s).

## Discussion

In this work, we present the development of a new optogenetic tool, optoSynC, for light-induced neuronal silencing *in vivo*. CRY2 homo-oligomerization, a modality thus far utilized mainly for induction of signaling pathways by protein-protein interaction^47,63,64^, was used to trigger formation of SV clusters, as evidenced by electron microscopy. This way, optoSynC activation efficiently and rapidly inhibits the release of neurotransmitters, as we showed by behavioral, pharmacological, and electrophysiological studies. optoSynC is, to our knowledge, the fastest acting and fully reversible optogenetic tool for inhibition of synaptic transmission that does not depend on membrane currents for hyperpolarization, or on induced protein damage.

CRY2 was targeted to the SV by fusion to SNG-1, an integral SV protein of high abundance, which increased the chance of efficient clustering of SVs. As SNG-1 is not required for synaptogenesis or neuronal activity^37^, optoSynC is inert in the dark, as we further showed by mutating the FAD-binding pocket of CRY2^32^. However, when activated with light, optoSynC causes neuronal inhibition within 25 s. Subsequently, in the dark, the effects recovered within 20 min. By specific expression, optoSynC can selectively interfere with activity of neuronal subtypes and even single neurons. By electron microscopy, we found clear evidence for SV approximation, as activation of optoSynC decreased the average distances of neighboring SVs by 14%. Had SV cluster formation occurred at the active zone, we would expect abnormal membrane structures, and defective SV recycling should lead to increased formation of large vesicles^53^. However, no such alterations were observed, although we noticed an accumulation of docked SVs right at the DP, while other docked SVs were depleted following optoSynC activation. Therefore, it seems most likely that SVs in the RP form clusters and cannot be transported to the active zone, thus causing a stop of transmission. Interaction of SVs through CRY2 oligomers seems not to prevent docking, yet those SVs may be inhibited in their mobility along the PM and accumulate at the DP. In optogenetically hyper-stimulated synapses, we previously observed that SVs became replenished along the PM, and only at late times, also re-appeared at the DP^53^. Thus, newly docked SVs (at the DP) diffuse laterally into the active zone membrane, until this pool is refilled, and only then start becoming visible at the DP. In optoSynC-stimulated synapses, the few free SVs that remain, may dock at the DP, but then be prevented from lateral diffusion by (hypothetical) optoSynC aggregates in the PM. In this context, the nearest distance of neighboring SV membranes in the RP, but not at the PM, was reduced by activated optoSynC from around 10 nm to 5 nm. The fact that we did not find zero distance of SVs in the RP may be due to the structural properties, and size, of CRY2^65^ and its oligomers, which likely limits the minimal distance of neighboring SVs.

To characterize the dynamics of optoSynC activation on synaptic transmission, we measured behavioral phenotypes. Activation of optoSynC impaired swimming behavior by 80%, which could be maintained for as long as illumination was applied (up to several hours, as shown in pharmacological assays). Impairment of swimming cycles occurred within seconds for some animals, and the full population effect developed over 25 s. However, optoSynC acts more rapidly, as we observed an immediate inhibition of photophobic escape behavior already during the first 2-3 s of illumination. By electrophysiology, the maximum effect took about one minute to develop. However, it is unknown how basal activity of the NMJ in dissected animals compares to activity during locomotion in an intact animal. In intact animals, inhibition of upstream interneuron activity increases the effects on behavior (compare pan-neuronal to cholinergic neuron-specific optoSynC). In dissected animals, interneuron - motor neuron connections may have been severed, thus possibly delaying optoSynC action.

optoSynC works with high performance: It mediates neuronal silencing at very low light intensities, swimming is decreased by 80%, mPSC frequency by 75%, and photophobic escape behavior is almost completely blocked. CRY2 clusters were described to be dynamic^45^, thus some transmitter may be released, possibly explaining remaining neuronal activity during blue light illumination. optoSynC is highly reversible, and swimming behavior recovered within 20 min in the dark. We recommend using an *sng-1* mutant background, since expression of optoSynC alongside endogenous SNG-1 reduced its efficacy, likely due to reduced incorporation into SVs. However, this depended on the neuron type and/or the promoter used, and did not affect efficiency in cholinergic neurons.

optoSynC is more efficient than other tools designed to inactivate neurotransmitter release by targeting SNARE proteins such as miniSOG/InSynC^25^, PSD^26,27^, or PA-BoNT^28^, particularly with respect to the fast action and recovery. These approaches affect protein degradation or inactivation, which is often quite slow, and is not reversible unless *de novo* synthesis occurs. miniSOG/InSynC^23^ reduces swimming behavior of *C. elegans*^26^, and can be used in single neurons; however, inhibition takes several minutes to build up and continues after the end of the light pulse. Inducing damaging radicals comes with off-target effects that can have unknown long-lasting outcomes^25^. PSD and PA-BoNT action (degradation/cleavage of SNT-1) reduce swimming behavior of *C. elegans* by 60-70 % (Refs. 26,28), but they require almost one hour. In comparison, optoSynC is more efficient, acts immediately, and can be sustained for several hours with no known off-target effects. Additionally, optoSynC reversibility is fast (20 min compared to 16-24 25,28), since no *de novo* synthesis of proteins is required. optoSynC has advantages also compared to light-driven hyperpolarizing ion pumps or anion channels^66^. Many light-driven ion pumps are only suitable for short-term silencing, as inactivation can reduce efficacy by about 50-90% within 60 s of illumination^19^. Light-gated anion channels rely on the chloride gradient, which makes use of these tools in neurons with unknown local chloride gradients problematic^14^, while in synaptic terminals they can actually induce depolarization^21^. optoSynC, however, acts completely independent of ionic gradients.

The recently developed opto-vTrap^36^, which is based on a LARIAT approach^35^, utilizes heterodimerization of CRY2 with CIB1^31,32^. We tried a similar approach expressing soluble CRY2 in synapses, and targeted CIBN to SV membranes *via* SNG-1. However, this did not reduce synaptic transmission. opto-vTrap targets VAMP, therefore SV fusion by blocking SNARE complex formation is the likely reason for inhibition, while optoSynC induces SV clusters. Thus, the tools are complementary; however, optoSynC may additionally enable investigation of different SV pools and individual steps during the SV cycle, as well as dynamics of SV mobilization. While opto-vTrap requires expression of two proteins, optoSynC is fully functional as a single tool. It should thus be easily transferable to other animal models, and the low light intensities needed for optoSynC likely facilitate applications further. optoSynC will be a useful tool for studies requiring precise, flexible, and reversible optogenetic silencing, as well as for studies of the dynamic interplay between SV pools.

## Materials and Methods

### Strains

*C. elegans* strains were cultivated according to standard methods^67^ on nematode growth medium (NGM) and fed *Escherichia coli* strain OP50-1. Animals were grown at room temperature and kept in the dark. Transgenic animals were obtained by microinjection of DNA into the gonads of animals in wt background (Bristol N2), *sng-1(ok234)*^37^ or *lite-1(ce314)* background^48^ by standard procedures^68,69^.

Strains used or generated: Bristol N2, *sng-1(ok234), lite-1(ce314)*, **ZX2577**: *sng-1(ok234); zxEx1216[psng-1::SNG-1::eGFP::CIBN; pmyo-2::mCherry]*, **ZX2581**: *sng-1(ok234); zxEx1224[psng-1::SNG-1::eGFP::CRY2olig(535); pmyo-2::mCherry]*, **ZX2604**: *sng-1(ok234); zxIs127[psng-1::SNG-1::CRY2olig(535);pmyo-2::mCherry]*, **ZX2628**: *sng-1(ok234); zxEx1234[psng-1::SNG-1::eGFP::CIBN; psng-1::mOrange2::CRY2olig(535); pmyo-2::mCherry]*, **ZX2737**: *zxIs132[punc-17::SNG-1::CRY2(535);pmyo-2::mCherry], ZX2807: zxIs137[ser2prom3::Chrimson::mNeonGreen; pmyo-2::mCherry]*, **ZX2816**: *zxEx1277[punc-47::SNG-1::CRY2olig(535); pmyo-3::mCherry]*, **ZX2865**: *zxIs137; zxEx1291 [ser2prom3::SNG-1::CRY2olig(535); pmyo-3::mCherry]*, **ZX2871**: *zxIs132; zxEx1277*, **ZX2872**: *lite-1(ce314); zxIs127*, **ZX2911**: *zxIs127*, **ZX2914**: *sng-1(ok234); zxIs132*, **ZX2950**: *sng-1(ok234)*; *zxEx1322[psng-1::SNG-1::CRY2(D387A)olig(535)::SL2::mCherry; pmyo-2::CFP]*.

### Molecular biology

To express optoSynC in *C. elegans*, the promoters p*sng-1* (pan-neuronal), p*unc-17* (cholinergic neurons), p*unc-47* (GABAergic neurons) and *ser2prom* (driving expression in the PVD neuron) were used. As selection marker, we expressed fluorescent proteins under the control of either the promoters p*myo-2* (expression in pharyngeal muscle) or p*myo-3* (expression in body wall muscle cells).

Plasmid **pDV01** [punc-17::CRY2olig(535)] was produced by amplifying the cDNA of CRY2olig(535) using primers oDV01 (5’-TGGCTAGCCGTCGTTCCGGAGGAGGTGGCGCCCGGGATCCAATGAAGATGGACAAAAAGACTATAGTTTGG -3’) and oMS67 (5’-CTGGGTCGAATTCGCCCTTTCCCTTGTCGACCATGACTCGAGTTAAACAGCCGAAGGTACTTGTTGG-3’), restriction digest of the fragment using *Sal*I and subcloning into vector pAH04 [punc-17::ccb-1::miniSOG::Citrine] by restriction digest with *Msc*I, *Psi*I and *Sal*I. Plasmid **pDV04** [punc-17::mOrange2::CRY2olig(535)] was generated by amplification of mOrange2 sequence with primers oDV013 (5’-CCGCATCTCTTGTTCAAGGGATTGGTGGCTAGGCTAGCCTCGAGAGGCCTGG-3’) and oDV014 (5’-GGGCGCCACCTCCTCCGGAACGACGGCTAGACTTGTAGAGTTCGTCCATTCCTCC-3’) and cloning it into vector pDV01 using *Nhe*I and Gibson Assembly. For **pDV09** [psng-1::mOrange2::CRY2(535)] the p*unc-17* sequence was replaced by the sequence of p*sng-1*, by restriction digest of pMS20 [psng-1::mCherry::BoNTB(N)::iLID] with *Msc*I and *Sph*I and pDV04 with *Nhe*I (blunted with T4 polymerase) and *Sph*I. For **pDV10** [psng-1::SNG-1::eGFP::CIBN], the CIBN fragment was amplified with primers oMS058 (5’-ATACAAAGGGGTTACCGGATCCGGCCTCGAGATGAATGGAGCTATAGGAGGTGAC-3’) and oMS059 (5’-TACGAATGCTCCCGGGCCTGCAGGCCCTAGGTTAAATATAATCCGTTTTCTCCAATTCCTTC-3’) and subcloning into pMS21 [psng-1::sng-1::eGFP::SspB(milli)_BoNTB(C)] using restriction enzymes *Avr*II and *Psp*XI. **pDV12** [psng-1::SNG-1::CRY2olig(535)] was assembled by amplification of CRY2olig(535) sequence with oMS060 (5’-GAGGGTCCGGTGGCGGAGGGTCAGGGGTACCGATGAAGATGGACAAAAAGACTATAGTTTG-3’) and oMS062 (5’-TACGAATGCTCCCGGGCCTGCAGGCCCTAGGTTATTTGCAACCATTTTTTCCCAAACTTG-3’) and subcloning into the psng-1::SNG-1 vector gained by restriction digest of pMS21 with *Avr*II and *Kpn*I. **pDV15** [psng-1::SNG-1::eGFP::CRY2olig(535)] was generated by amplifying the CRY2olig(535) fragment with oMS062 (5’-TACGAATGCTCCCGGGCCTGCAGGCCCTAGGTTATTTGCAACCATTTTTTCCCAAACTTG-3’) and oMS063 (5’-ATACAAAGGGGTTACCGGATCCGGCCTCGAGATGAAGATGGACAAAAAGACTATAGTTTG-3’) and Gibson assembled with the psng-1::SNG-1::eGFP fragment after restriction digest of pMS21 with *Avr*II and *Psp*XI. The plasmid **pDV06** [punc-17::SNG-1::CRY2(535)] was generated by restriction digest of pDV12 and pMSM17 [5’hom-CCB-1::mKate2::SEC::cMyc(3x)::iLIDpsd-s::3’hom-CCB-1] with *Kpn*I and *Bsm*I and subsequent ligation. For **pDV18** [punc-47::SNG-1::CRY2olig(535)], p*unc-17* was replaced by amplification of the p*unc-47* promoter with oDV025 (5’-AACAACTTGGAAATGAAATAAGCTTGCATGCCTGCAGAGCTTGTTGTCAT-3’) and oDV026 (5’-GCACCATAAGCACGCACGTTCTCCATTTCACCGGTGCTGTAATGAAATAAATGTGACGC-3’) and subcloning into restriction-digested pDV06 with *Sph*I-HF and *Sgr*AI-HF. Plasmid **pDV19** [ser2prom3::SNG-1::CRY2olig(535)] was generated by ligation of the restriction digest of pDV06 and pJW41 [ser2prom3::Chrimson::mNeon] with *Bmt*I-HF (blunted with T4 polymerase) and *Sph*I-HF. To construct **pDV20** [pSNG-1::SNG-1::CRY2(D387A)olig(535)], site-directed mutagenesis was conducted using the template pDV12 and the primers oDV027 (5’-ACACTTTTGGCTGCTGATTTGG-3’) and pDV028 (5’-CCAAATCAGCAGCCAAAAGTGT-3’). **pDV21** [pSNG-1::SNG-1::CRY2(D387A)olig(535)::SL2::mCherry] was generated by ligation of the restriction digests of pDV20 with *Pvu*I-HF and *Sma*I and pTH02 [pmyo-3::CaCyclOp::SL2::mCherry] with *Kpn*I-HF (blunted using T4 polymerase) and *Pvu*I-HF.

### Behavioral assays

All strains were kept in the dark on standard NGM plates (6 cm, 8 ml NGM) with OP50-1 bacteria at room temperature. For analysis of swimming behavior, transgenic L4 larvae were selected for fluorescent markers under a Leica MZ16F dissection scope and transferred to freshly seeded NGM plates. After 12-16 h in the dark, 60-80 young adult animals were transferred onto plain NGM plates (3.5 cm diameter, 3 ml NGM) using 800 µl M9 buffer. Animals were kept in darkness for 15 min for accommodation to swimming behavior. Then, the plate was placed onto a multiworm tracker (MWT) platform^70^ equipped with a high-resolution camera (Falcon 4M30, DALSA) and a custom-built infrared transmission light source (6 WEPIR3-S1, WINGER, 850nm 3W). Using a LabVIEW-based custom software (MS-Acqu), videos of swimming behavior were automatically recorded for 1 min, at different time points during the experiment. A custom-built LED ring (Alustar 3W 30°, ledxon, 470 nm) was used for light stimulation controlled by transistor-transistor logic (TTL) pulses from MS-Acqu. Afterward, swimming cycles were automatically analyzed using “wrMTrck” plugin^71^ for ImageJ (version 1.52a, National Institute of Health). Tracking was validated, non-worm objects manually removed, and data summarized using custom Java and Python scripts. For the acquisition of speed data, the crawling behavior of young adult animals was analyzed on the MWT. Worms were collected in M9 buffer, transferred in a droplet of buffer onto a fresh plain NGM plate (6 cm, 8 ml NGM), and kept for 15 min in the dark. Speed data was recorded as described^70^. Tracks were extracted using Choreography and output files organized into summary statistics using a custom Python script.

### Pharmacological assays

To assay aldicarb sensitivity, 2 mM aldicarb plates were prepared^51^. After cultivation in the dark at room temperature, animals were transferred to the aldicarb dishes (15-27 young adults per trial, three biological replicates in total on three consecutive days, animals picked from different populations) and scored every 30 min, for up to 6 h, by three gentle touches with a hair pick to nose and tail regions. All conditions were recorded blinded on the same day. For analysis of the effect of activated optoSynC, animals were illuminated with blue light (470nm, 0.05 mW/mm ^2^) throughout the 6 h experiment.

### Electrophysiology

Electrophysiological recordings from body wall muscle cells were done in dissected adult worms as described^72^. Animals were immobilized with Histoacryl glue (B. Braun Surgical, Spain) and a lateral incision was made to access NMJs along the anterior ventral nerve cord. Removal of the basement membrane overlying body wall muscles was enzymatically achieved by incubation in 0.5 mg/ml collagenase for 10 s (C5138, Sigma Aldrich, Germany). The integrity of body wall muscle cells and nerve cord was examined via DIC microscopy. Recordings from BWMs were acquired in whole-cell patch-clamp mode at 20-22°C using an EPC-10 amplifier equipped with Patchmaster software (HEKA, Germany). The head stage was connected to a standard HEKA pipette holder for fire-polished borosilicate pipettes (1B100F-4, Worcester Polytechnic Institute, USA) of 4-10 MΩ resistance. The extracellular bath solution consisted of 150 mm NaCl, 5 mm KCl, 5 mm CaCl2, 1 mm MgCl2, 10 mm glucose, 5 mm sucrose, and 15 mm HEPES, pH 7.3, with NaOH, ∼330 mOsm. The internal/patch pipette solution consisted of K-gluconate 115 mm, KCl 25 mm, CaCl2 0.1 mm, MgCl2 5 mm, BAPTA 1 mm, HEPES 10 mm, Na2ATP 5 mm, Na2GTP 0.5 mm, cAMP 0.5 mm, and cGMP 0.5 mm, pH 7.2, with KOH, ∼320 mOsm. Activation of light was performed using a LED lamp (470 nm, 8 mW/mm^2^; KSL-70, Rapp OptoElectronic, Germany), controlled by an EPC-10 amplifier and Patchmaster software (HEKA, Germany). Samples were illuminated using a 5 s/5 s ISI for 30 s. Analysis of mPSCs was done using the Mini Analysis software (Synaptosoft). Amplitude and mean number of mPSC events per second were analyzed during the following time bins: 30 s before illumination and 20 s to 30 s after illumination.

### Electron Microscopy

High-pressure freezing (HPF) fixation of young animals was performed as described earlier^53,73^. In brief, 20-40 worms were transferred into a 100 µm deep aluminum planchette (Microscopy Services) filled with *E. coli* OP50, covered by a 0.16 mm sapphire disk and a 0.4 mm spacer ring (Engineering office M. Wohlwend) for photostimulation. To prevent the preactivation of optoSynC, handling of worms was performed under red light. Animals were illuminated with a 470 nm blue LED (0.1 mW/mm^2^) for 5 s followed by high-pressure freezing after 25 s at -180°C under 2100 bar pressure in an HPM100 machine (Leica Microsystems). After freezing, samples were transferred under liquid nitrogen into a Reichert AFS machine (Leica Microsystems) for freeze-substitution. Samples were incubated in tannic acid (0.1% in dry acetone) fixative at -90°C for 100 h. This was followed by a process of washing to substitute with acetone and incubation in 2% OsO4 for 39.5 h (in dry acetone) while slowly increasing the temperature up to room temperature. Afterwards, an epoxy resin (Agar Scientific, AGAR 100 Premix kit hard) embedding process was performed with increasing concentration from 50% to 90% at room temperature and 100% at 60°C over 48 h. Cross-sections were cut at a thickness of 40 nm, transferred on formvar-covered copper slot grids, and counterstained with 2.5% aqueous uranyl acetate for 4 min, followed by washing steps with distilled water. Then, grids were incubated in Reynolds lead citrate solution for 2 min in a CO2-free chamber and washed again with distilled water. The ventral nerve cord region was imaged using a Zeiss 900 TEM, operated at 80 kV, and a Troendle 2K camera. For analysis of images, plasma membrane, dense projection, synaptic vesicles (SVs), docked SVs, dense core vesicles (DCVs) and large vesicles (LVs) were annotated in ImageJ (version 1.53c, National Institute of Health) using the synapsEM workflow^57^. Annotated images were analyzed using affiliated MATLAB (R2021a, MathWorks) scripts and a custom script calculating nearest vesicle distances.

### Data and statistical analysis

Data are shown as mean ± s.e.m. or as median with interquartile range, n indicates the number of animals, and N the number of biological replicates. Significance between datasets after one- or two-way ANOVA with Bonferroni’s multiple comparison test is given as a p-value. If data without normal distribution were compared, we used Mann-Whitney’s test. Comparison between relative frequency distributions was compared with Kolmogorov-Smirnov test. The respective statistics used are indicated in the figure legends. Data were analyzed and plotted in GraphPad Prism (GraphPad Software, version 8.02).

## Author Contributions

Conceptualization: MS, DV, AG

Methodology / Software: SW, MS, DV

Investigation: DV, MS, JFL, SZ, JG

Visualization: DV, AG

Funding Acquisition: AG

Project administration: AG

Supervision: AG

Writing – original draft: DV

Writing – review & editing: AG, DV, SW, JFL, JG

## Acknowledgements

We thank Chandra Tucker for providing CRY2 plasmids. We thank members of the Gottschalk lab for critical comments. We acknowledge the *Caenorhabditis* Genetic Center (CGC), which is funded by NIH Office of Research Infrastructure Programs (P40 OD010440), and the National Bioresource project, nematode *C. elegans*, for strains. We are indebted to Franziska Baumbach, Hans-Werner Müller, Barbara Janosi, Marion Basoglu, and Marius Seidenthal for expert technical assistance. This work was supported by grants from the Deutsche Forschungsgemeinschaft (DFG), SPP 1926, Project VIb (GO1011/12-2), and by Goethe University Frankfurt to AG. SW is supported by National Institutes of Health (1DP2 NS111133-01 and 1R01 NS105810-01A1. SW is an Alfred P. Sloan fellow, McKnight Foundation Scholar, and Klingenstein and Simons Foundation scholar.

## Supplementary Materials

### Supplementary Figures

**Supplementary Figure S1.**
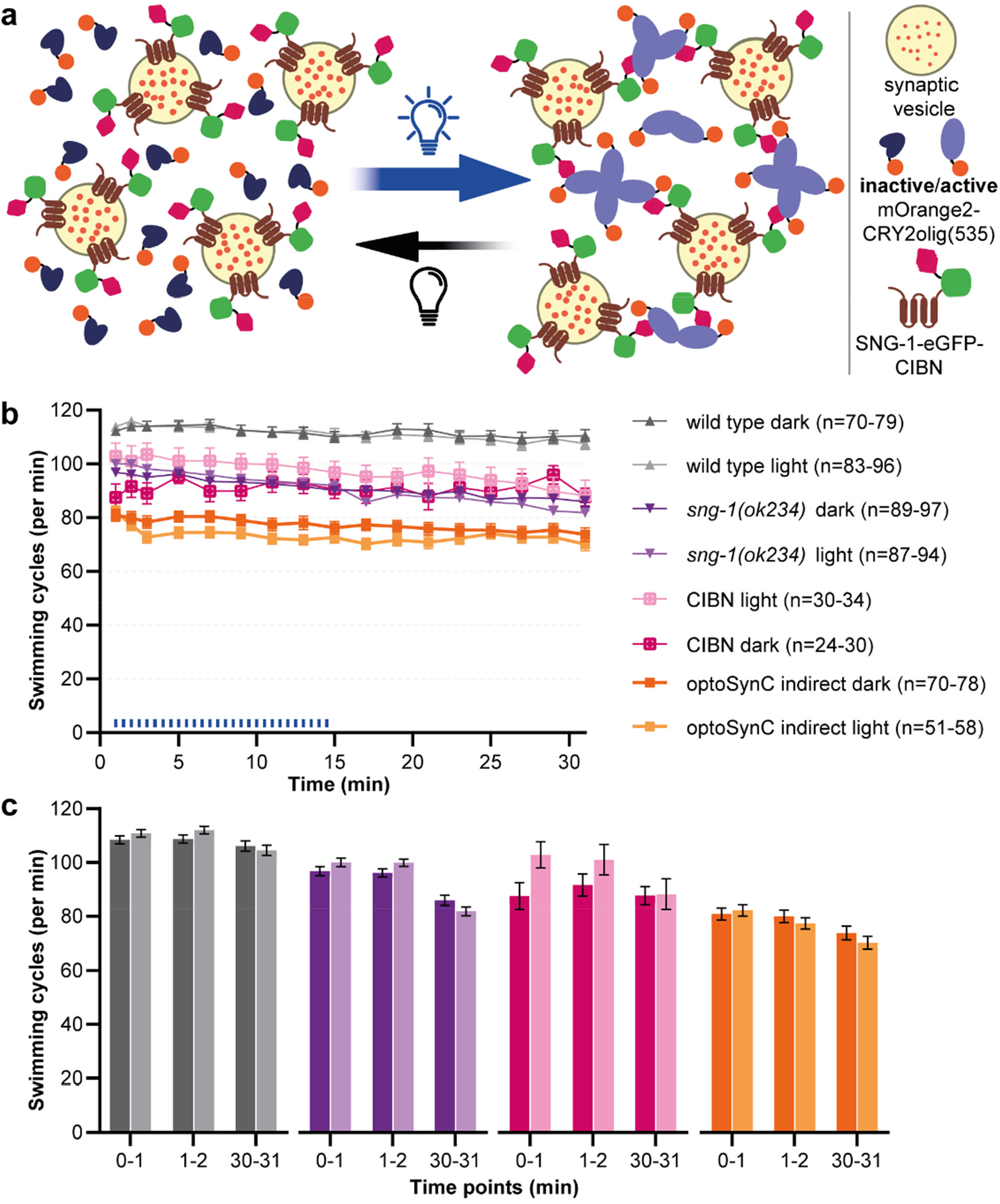
LARIAT-based approach did not affect swimming behavior: **a** Schematic of LARIAT-based approach (‘optoSynC indirect’) utilizing heterodimerization of SV membrane bound CIBN and cytosolically expressed CRY2olig(535). **b** Mean ± s.e.m. swimming behavior (cycles) of animals expressing the indicated proteins. Blue light pulses (470 nm, 0.1 mW/mm^2^, 5 s / 25 s ISI) are indicated by blue bars. **c** Analysis of data in (b) during intervals before (0-1 min), during (1-2 min) and after (30-31 min) blue light illumination. Two-way ANOVA with Bonferroni correction between light and dark; *p<0.05, **p<0.01, ***p<0.001. Strains as indicated in (b). From left to right, number of animals n=107, 120, 104, 120, 93, 115, 96, 87, 95, 88, 89, 92, 98, 26, 34, 24, 33, 26, 31, 72, 58, 74, 57, 71 and 54.

**Supplementary Figure S2.**
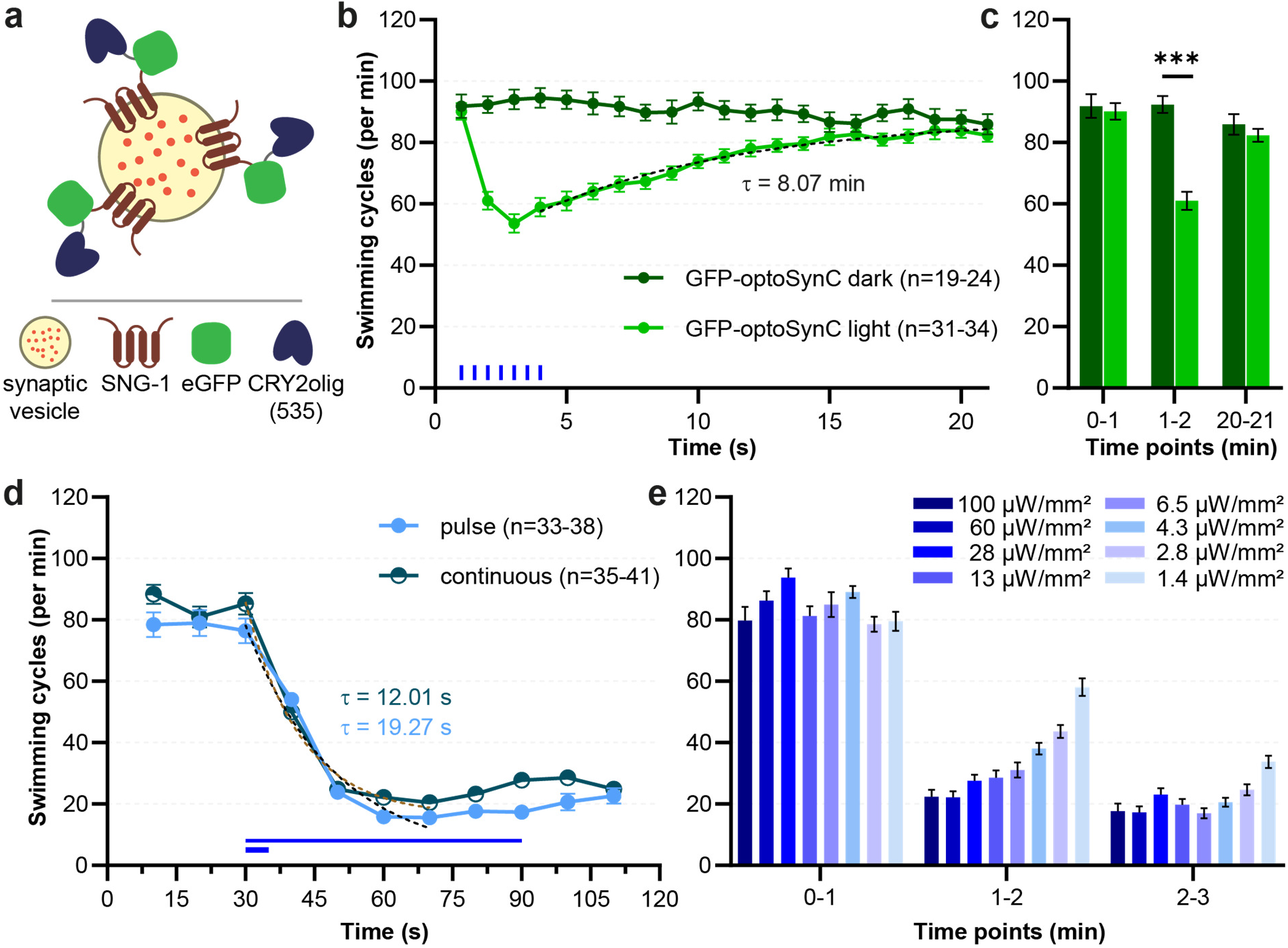
Inserting a fluorescent protein in optoSynC reduces activity; required pulse duration and light sensitivity of optoSynC: **a** Schematic illustration of GFP-optoSynC construct. eGFP is inserted between SNG-1 and CRY2olig(535). **b** Mean ± s.e.m. of swimming behavior before, during and after activation of GFP-optoSynC with blue light (470nm, 0.1 mW/mm^2^, 5 s / 25 s ISI, blue rectangles). Dotted line represents one-phase association fit during recovery of swimming behavior. **c** Mean data in (b), analyzed in intervals before (0-1 min), during (1-2 min) and after (20-21 min) blue light illumination. Two-way ANOVA with Bonferroni correction; *p<0.05, **p<0.01, ***p<0.001. From left to right, number of animals is n=23, 33, 24, 34, 23 and 32. **d** OptoSynC can be fully activated by a single short pulse of blue light (470 nm, 0.1 mW/mm^2^, 5 s); compare to continuous 1 min illumination. Dotted lines represent one-phase decay after pulsed or continuous illumination. **e** Light sensitivity of optoSynC, tested by swimming behavior (mean ± s.e.m.). The indicated intensities of illumination were used during 3 min experiments. Analysis during the indicated intervals.

**Supplementary Figure 3.**
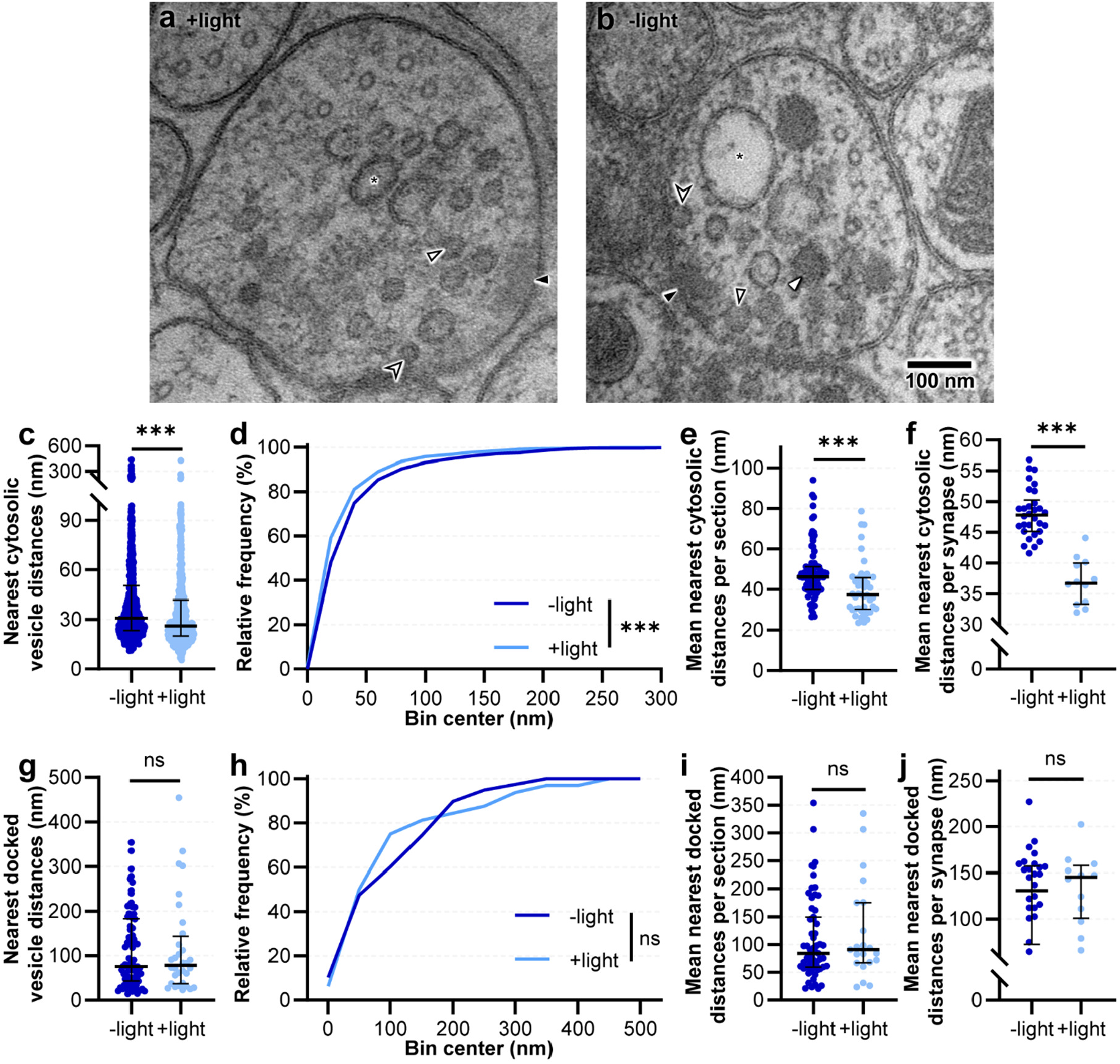
Ultrastructural parameters of cholinergic terminals following optoSynC activation: **a, b** Transmission electron micrographs of representative cholinergic synapses from animals illuminated for 5 s (a, +light) with blue light (470 nm, 0.1 mW/mm^2^) or kept in darkness (b, -light) before high-pressure freezing. Synaptic vesicles (SVs, open black arrowheads), dense core vesicles (DCVs, white closed arrow heads), dense projection (DP, closed black arrowheads), docked SVs (open black notched arrowhead) and large vesicles (LVs, asterisks) are indicated. **c** Distance analysis of cytosolic vesicles for each analyzed cholinergic micrograph shown as median with interquartile range. Mann-Whitney test between -light (n=1327) and +light (n=755). **d** Relative frequency distribution of nearest cytosolic vesicle distances shown in (c). Kolmogorov-Smirnov test. **e** Mean nearest cytosolic distances per section shown as median with interquartile range. Mann-Whitney test between -light (n=88) and +light (n=43). **f** Mean nearest distances per synapse shown as median with interquartile range. Mann-Whitney test between -light (n=30) and +light (n=12). **g** Distance analysis of docked vesicles for each analyzed cholinergic micrograph shown as median with interquartile range. Mann-Whitney test between -light (n=78) and +light (n=32). **h** Relative frequency distribution of nearest docked vesicle distances shown in (g). Kolmogorov-Smirnov test. **i** Mean nearest docked distances per section shown as median with interquartile range. Mann-Whitney test between -light (n=59) and +light (n=21). **j** Mean nearest docked distances per synapse shown as median with interquartile range (Mann-Whitney test between -light (n=30) and +light (n=12). Statistically significant differences in (c-j) are given by ***p<0.001, ns not significant.

**Supplementary Figure 4.**
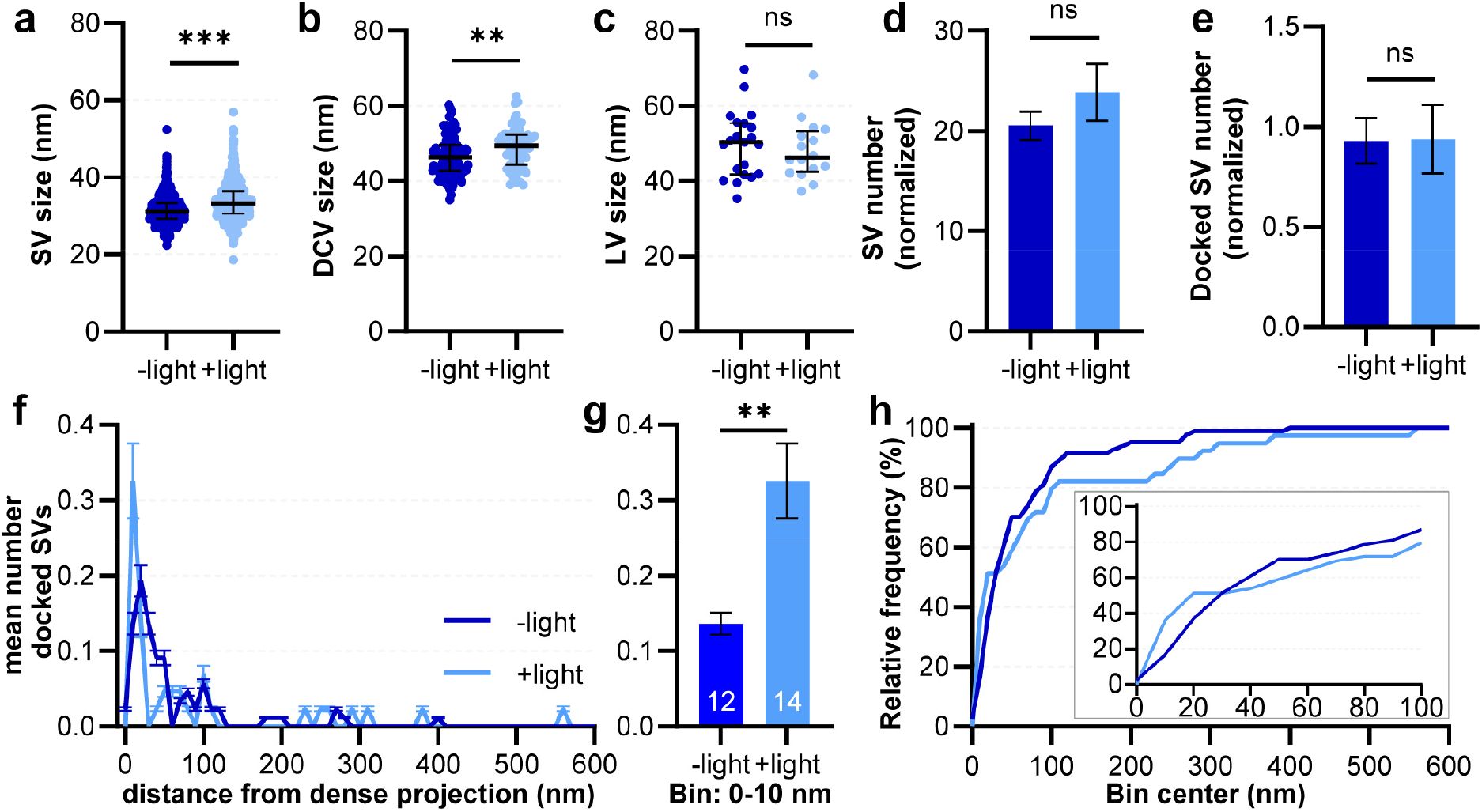
Vesicle numbers and distribution in cholinergic terminals following optoSynC activation: **a-c** Analysis of diameters of SVs (a, n=1561 and 862, from left to right), DCVs (b, n=133 and 56) and LVs (c, n=21 and 16) shown as median with interquartile range (Mann-Whitney test). **d, e** Normalized number of SVs (d) and docked SVs (e) by multiplying with length of plasma membrane of each analyzed micrograph and dividing through mean length of PM of each data set (Error bars are s.e.m., Mann-Whitney test between -light (n=88) and +light (n=43). **f** Distribution of distances of docked SVs relative to the DP, along the PM, in 10 nm bins. **g** Number of docked SVs in bin 1-10 nm, i.e. directly neighboring the DP. **h** Relative frequency distribution of docked vesicle distances to the DP, shown in (f). Kolmogorov-Smirnov test: n.s. Inset shows enlarged region of bins 10-100 nm. Statistically significant differences in (a-e, g) are given by **p<0.01, ***p<0.001, ns not significant.

**Supplementary Figure 5.**
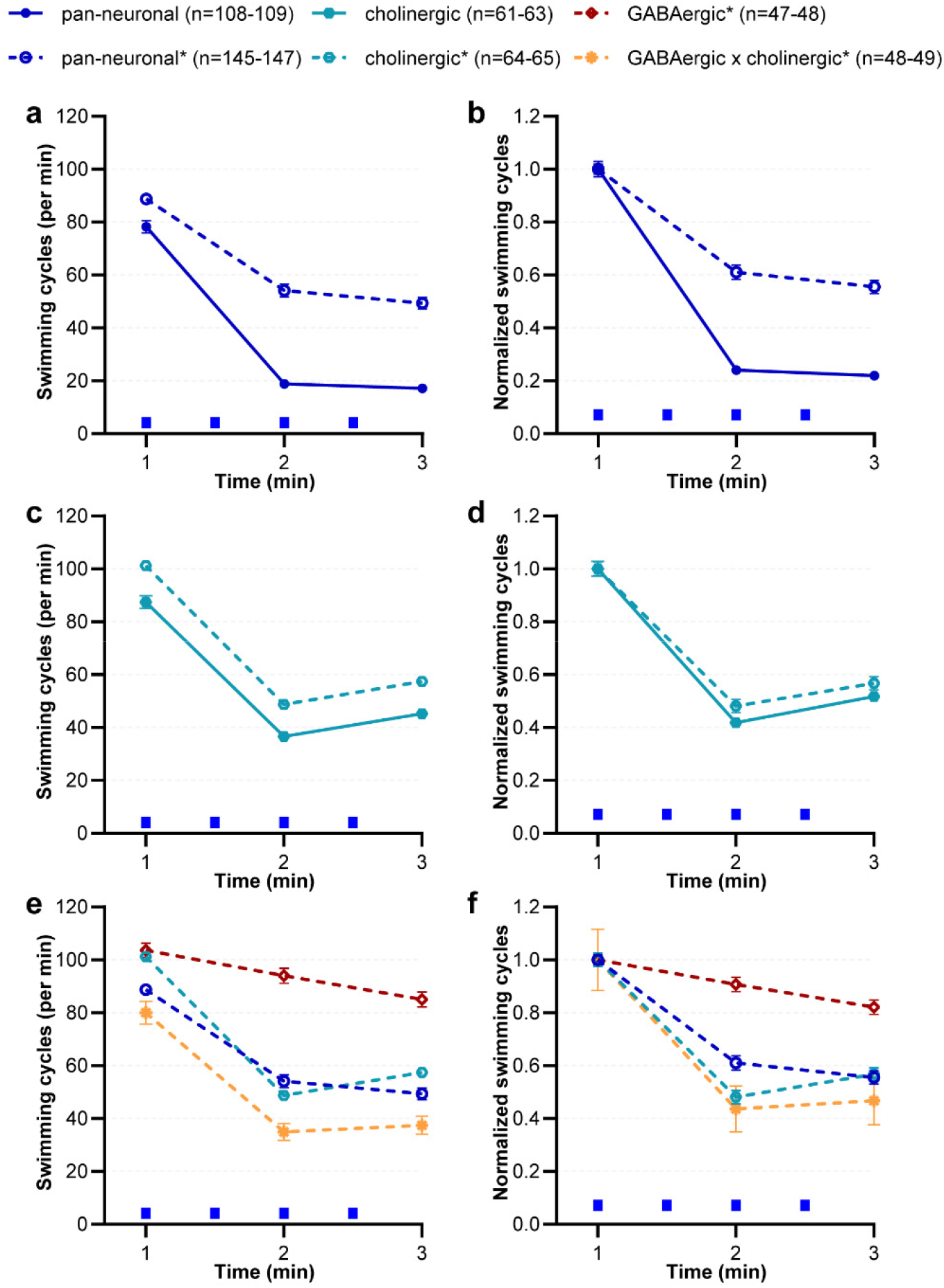
Competition between optoSynC and endogenous SNG-1 can reduce optoSynC effects, dependent on neuron type: **a, b** Swimming behavior (normalized in b) before and after activation by blue light (470 nm, 0.1 mW/mm ^2^, 5 s / 25 s ISI) was analyzed in animals expressing optoSynC pan-neuronally in wild type worms (indicated by asterisk in figure legend and dotted line) vs. *sng-1(ok234)* mutants. **c, d** As in (a, b), but comparing animals expressing optoSynC in cholinergic neurons, in wild type and *sng-1(ok234)* mutants. **e, f** Comparing swimming behavior in animals expressing optoSynC in GABAergic or cholinergic neurons, in both cell types, or panneuronally, in wild type background.

### Supplementary Video

**Supplementary Video 1: Rapid inhibition of swimming locomotion after optoSynC pan-neuronal photoactivation**. Video speed is increased 2x. Blue rectangle indicates application of light pulse. Time is shown as min:sec:centisec.

